# Detection of viral infection in cell lines using ViralCellDetector

**DOI:** 10.1101/2023.07.21.550094

**Authors:** Rama Shankar, Shreya Paithankar, Suchir Gupta, Bin Chen

## Abstract

Cell lines are commonly used in research to study biology, including gene expression regulation, cancer progression, and drug responses. However, cross-contaminations with bacteria, mycoplasma, and viruses are common issues in cell line experiments. Detection of bacteria and mycoplasma infections in cell lines is relatively easy but identifying viral infections in cell lines is difficult. Currently, there are no established methods or tools available for detecting viral infections in cell lines. To address this challenge, we developed a tool called ViralCellDetector that detects viruses through mapping RNA-seq data to a library of virus genome. Using this tool, we observed that around 10% of experiments with the MCF7 cell line were likely infected with viruses. Furthermore, to facilitate the detection of samples with unknown sources of viral infection, we identified the differentially expressed genes involved in viral infection from two different cell lines and used these genes in a machine learning approach to classify infected samples based on the host response gene expression biomarkers. Our model reclassifies the infected and non-infected samples with an AUC of 0.91 and an accuracy of 0.93. Overall, our mapping- and marker-based approaches can detect viral infections in any cell line simply based on readily accessible RNA-seq data, allowing researchers to avoid the use of unintentionally infected cell lines in their studies.

## INTRODUCTION

Cell lines are a valuable tool in clinical research as they allow scientists to study disease mechanisms and develop new treatments. They can be used to study a wide range of biological processes, including cell signaling, gene expression, and drug metabolism. Additionally, cell lines are widely used to test the efficacy and toxicity of new drugs before in vivo evaluation. Cell lines can be derived from a variety of sources, including patient cancer cells, human primary cells, or cells from animals. Immortalized cancer cell lines are often shared across labs and studies. However, cross-contamination with bacteria and mycoplasma is a significant issue (Corral-Vázquez *et al*., 2017). To avoid these contamination issues, best practice guidelines should be followed (Baust *et al*., 2017; Reid, 2017).

In addition to bacterial and mycoplasma contamination, viral contamination is also a major constraint in cell lines, yet difficult to detect. Viral contamination can come from the environment or from the tissue from which the cell lines were derived. Although human cell lines may potentially carry a virus, there are limited approaches for detecting these viruses in cell lines (Shioda *et al*., 2018; Cheval *et al*., 2019; Uphoff *et al*., 2019; Dolskiy *et al*., 2020). These approaches are limited to a few viruses such as cytomegalovirus (CMV), Epstein-Barr virus (EBV), human herpesvirus 6 (HHV-6), HHV-7, human polyomavirus BK (BKV), human polyomavirus JC (JCV), human adenovirus (ADV), human parvovirus B19 (B19V), hepatitis B virus (HBV), human T-cell leukemia virus type 1 (HTLV-1), HTLV-2, human immunodeficiency virus 1 (HIV-1), HIV-2, hepatitis A virus (HAV), and hepatitis C virus (HCV), which are known to be pathogenic. The methods based on either PCR or specific viral sequence reads have to be tailed for specific viruses, thus challenging to generalize to research.

Recently, several algorithms have been developed to detect virus integration in the host genome based on sequencing data (Kostic *et al*., 2011; Y. Chen *et al*., 2013; Li *et al*., 2013; Naeem *et al*., 2013; Schelhorn *et al*., 2013; Wang *et al*., 2013, 2015; Xia *et al*., 2019; Selitsky *et al*., 2020). Among them, VirTect (Xia *et al*., 2019) is the most advanced and can detect virus integration sites based on whole transcriptome sequencing (RNA-Seq) data. However, there is no software or tool is available to detect the viral infection in any sample. The mapping-based approaches often posit that viral sequence reads are retained in the cell line sequence data; however, the widely used polyA based library preparation protocol only include the reads from the host with poly-adenylated. Therefore, there is a need for a new tool to detect viruses in any samples using RNA-seq data.

To overcome these limitations, we first developed a new tool that utilizes the ultrafast STAR aligner (Dobin *et al*., 2013) followed by BWA aligner and includes all the viruses present in the NCBI virus database. Our tool begins by mapping RNA-seq data to the human genome and transcriptome, followed by mapping unmapped reads to a viral genome database. We removed the viruses that belong to plants or invertebrates and apply stringent criteria to accurately detect true viruses. Furthermore, we implemented a machine learning approach to identify biomarkers for detecting viral infections in samples. This biomarker-based approach has remarkable performance and is independent from virus types and library preparation protocol; however, it is only appropriate in the samples where viral infection changed the biology of the host cells. We expect that the combination of both mapping and biomarker-based approach will aid researchers in detecting suspicious viral infections in their cell lines using RNA-seq data and control the quality of their experiments.

## METHODS

The overall workflow of ViralCellDetector is shown in Fig. 1A. ViralCellDetector takes as input the fastq files of paired-end sequencing reads from each sample. The first step of ViralCellDetector is to align the sequencing reads to the Hg38 transcriptome with ENSEMBL GRCh38.p3 annotation using STAR aligner (Dobin *et al*., 2013). Unmapped reads are then further mapped to the viral genome database available on NCBI (https://ftp.ncbi.nlm.nih.gov/refseq/release/viral/) using BWA (Li and Durbin, 2009). The advantage of using BWA over STAR aligner for viral genomes is that it can reports mapping of reads even if only one read from paired read is mapped on the viral genome. The total number of mapped reads on the viral genome is calculated to remove false-positive viruses. Additionally, the percentage coverage of viral genomes is estimated to identify true viruses. It is important to note that individual samples’ sequencing reads should be mapped individually on the human transcriptome and viral genome.

**Fig. 1.**
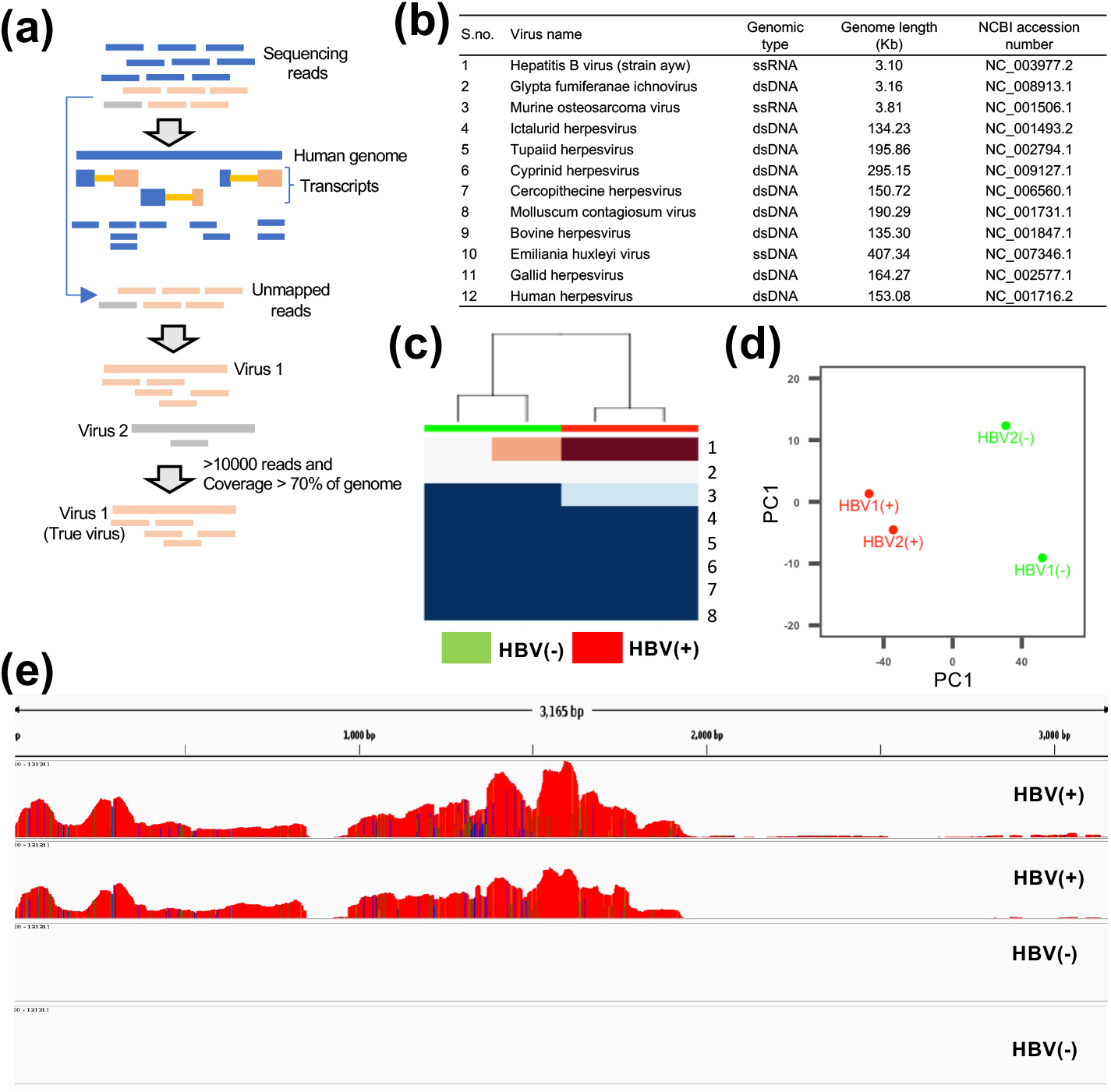
Detection of viral infection using VirusCellDetector. (a) The workflow of VirusCellDetector for viral detection. (b) A list of viruses detected in HBV-positive (HBV+) and control (HBV-) samples. (c) Heatmaps illustrating the normalized distribution of viruses in HBV+ and HBV-samples. (d) Principal component analysis of HBV+ and HBV-samples. (e) Integrated Genomics Viewer displays of base coverage for the HBV genome in HBV+ and HBV-samples. Multiple tracks for four libraries show the reads mapped to each location, based on alignments to the hepatitis B virus reference genome.

### Dataset

We downloaded the SRA sequencing data from GEO (https://www.ncbi.nlm.nih.gov/geo/browse/?view=series). All the sequencing data were further converted into fastq files using the SRAtoolkit (https://github.com/ncbi/sra-tools/wiki/01.-Downloading-SRA-Toolkit). The fastq files were used as input in the ViralCellDetector to detect the putative viruses.

### Data processing

ViralCellDetector is designed to accept raw sequencing data as input. ViralCellDetector separates the paired end reads into two sets: those that belong to the reference genome, and those that belong to viral genomes. The reads that belong to the host reference were mapped to the reference genome and transcriptome using the ultrafast STAR aligner with default parameters (Dobin *et al*., 2013). The unmapped reads left after host reference mapping were mapped to the NCBI viral genome database using the BWA aligner with default parameters (Li and Durbin, 2009). The viral genome sequences can be downloaded from the NCBI virus database (https://ftp.ncbi.nlm.nih.gov/refseq/release/viral/). Once the viral reads are mapped to the viral genome, several criteria were applied to filter out false positives, such as: 1) at least 10,000 reads should be mapped to the viral genome, 2) the continuous coverage of the viral genome should be more than 70%, and 3) the virus should be known to infect vertebrates.

### RNA-seq analysis

All the sequencing reads were mapped on Hg38 transcriptome using the ENSEMBL GRCh38.p3 annotation with the STAR aligner (Dobin *et al*., 2013). The gene count obtained after mapping was used for identification of differentially expressed (DE) genes. The edgeR package (Robinson *et al*., 2010) was used for quantification of DE genes with criteria: log2 fold change ≥1 or ≤-1 with adjusted P-value (False Discover Rate) ≤ 0.01. DE genes were identified between the two groups in two different independent datasets. we performed gene ontology enrichment and KEGG pathways analyses using the enrichR (E. Y. Chen *et al*., 2013; Kuleshov *et al*., 2016). DE genes involved in viral infection related pathways were used further as feature in machine learning. The data was visualized using the ggplot2 package in R. R 3.5.1 was used to perform the different analyses.

### Random forest for feature selection and prediction

Random forest (RF) is a robust machine learning algorithm that is based on bagging techniques and consistently outperforms others in various machine learning tasks (Subash *et al*., 2022; Wadood *et al*., 2022). It consists of multiple decision trees that are independent of each other, and the final output of the model is determined by aggregating the results from each decision tree in the forest (Montes *et al*., 2021). The three parameters that affect the performance of the model are “ntree,” “mtry,” and “nodesize” and can be tuned by the user (Oukawa *et al*., 2022). We used the ntree value 600 and mtry 8 parameters in the R random forest classifier for feature selection and prediction. The experimental data was divided into 80% training set, and the remaining 20% was used as a test dataset. Recursive feature selection method was used for feature selection. To evaluate the robustness of the predictive model for feature selection, 10-fold cross-validation was applied. Features with an accuracy of ≥ 0.9 in 10-fold cross-validation were selected for prediction in the test dataset. Finally, accuracy, AUC, sensitivity, and specificity were calculated for the test dataset.

## RESULTS

### Validation of the mapping-based viral infection detection approach

The workflow of the pipeline is provided in Fig.1a. We collected datasets with known infected samples. One dataset included Hepatitis B virus (HBV) infected samples (two control and two infected) (GSE65485), and the other dataset belonged to SARS-CoV-2 infected cell lines (three control and three infected) (GSE187420). The HBV dataset included two control samples and two HBV infected samples. In this dataset, we obtained a total of 12 viruses in the list (Fig. 1b). However, when we checked for expression patterns in the control and infected samples, we observed that the HBV infected samples showed the highest expression levels as compared to the control samples (Fig. 1c). Additionally, these two groups of samples were transcriptionally different from each other (Fig. 1d). Finally, when we visualized the reads mapped to the different viral genomes, we observed that only HBV genome is covered more than 90% with the reads (Fig. 1e). This confirms the HBV infection in the sample. Additionally, the other dataset (GSE187420) confirmed a high unmapped read count on host reference transcriptome (Table S1) and very high SARS-CoV-2 read count in the SARS-CoV-2 infected samples (Fig. S1). The two cases validated the approach used in this method for detecting viral infections in cell lines.

### Detection of viral infection from unlabeled dataset

To detect viral infections in unlabeled (without known infection) RNA-seq data, we collected MCF7 cell line datasets from multiple studies (SRP065220, SRP142602, and SRP163132). The SRP065220 dataset includes six samples (three controls and three decitabine treatments), the SRP142602 study includes four MCF7 samples, and SRP163132 has 128 samples. The sequencing data in these studies were generated by a RiboMinus approach, which enables to capture all expressed mRNAs from human and other species including virus (Zhao *et al*., 2018; Chen *et al*., 2020). The raw reads were downloaded and first mapped to the human transcriptome. Then the total unmapped reads were computed for each sample. We observed that eight samples had more than 45% unmapped reads (Fig. 2a) suggesting that these samples might have viral infections, as it has been observed that samples with viral infections have high unmapped reads as compared to controls in our data (Table S1) as well as previous report (Yuan *et al*., 2021). The unmapped reads were then mapped to the viral genome using ViralCellDetector and identified a total of 22 viruses (Fig. S2). Further by applying stringent cutoffs, we were able to identify a total of six viruses containing three dsDNA viruses (Woodchuck hepatitis virus, BeAN 58058 virus, and Eptesicus fuscus gamma herpes virus) and three ssRNA viruses (Human immunodeficiency virus 1, Encephalomycarditis virus and Human endogenous retrovirus) (Fig. 2b).

**Fig. 2.**
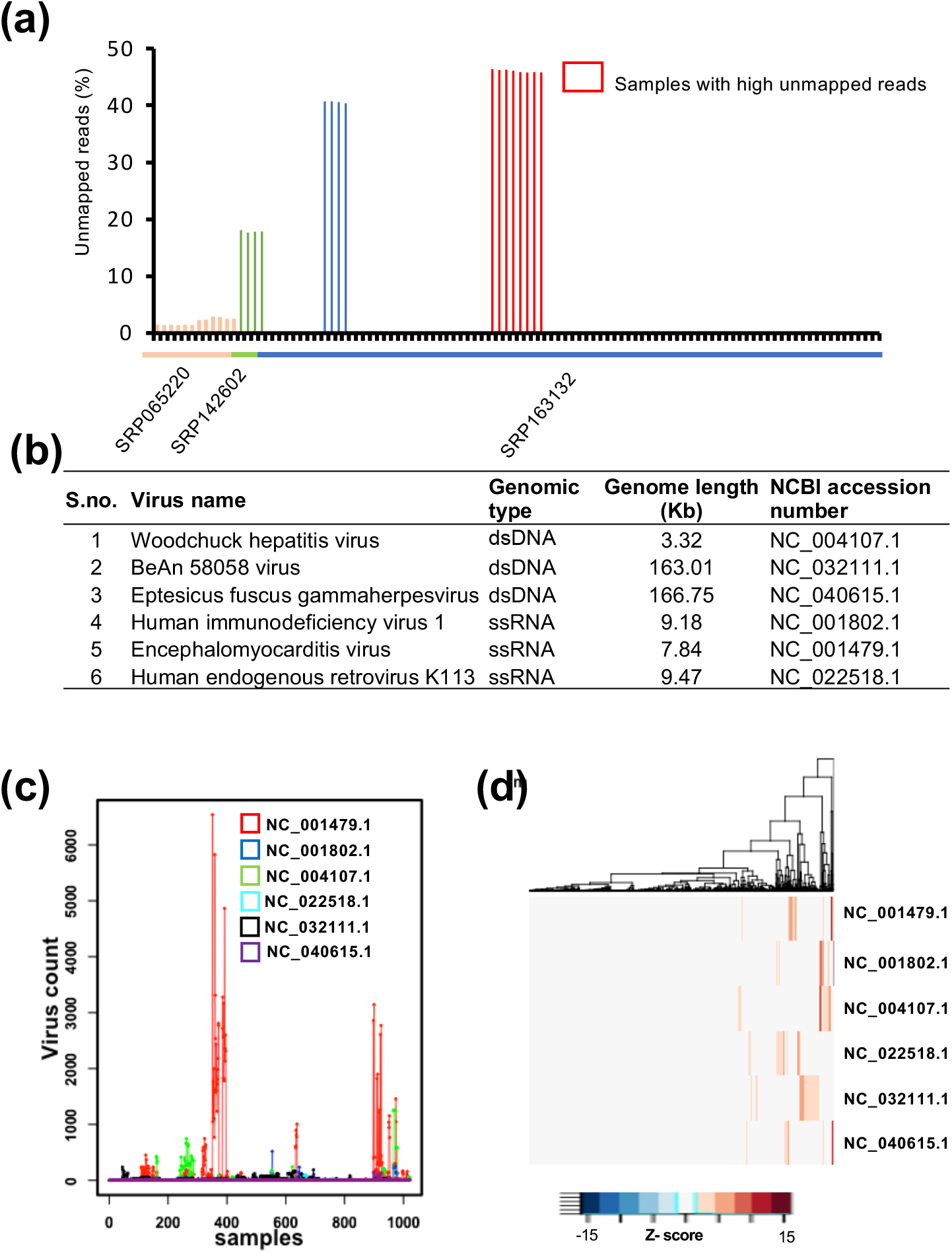
Detection of viral infection using VirusCellDetector on experimental data of the MCF7 cell line. (a) Percentage of unmapped reads in different samples from three different studies. The highest percentage of unmapped reads was 45% in eight samples from the SRP163132 study. These samples were used for viral detection and resulted in the identification of a total of six viruses. (b) The expression of these six viruses was analyzed in 1018 experiments of MCF cells, and many samples were found to be infected (as shown in Fig. c). (d) Heatmaps illustrating the higher expression of the six viruses in various experiments in MCF7 cells.

To further explore the presence of these viruses in MCF7 cells, we collected RNA-seq data from over 1000 MCF7 samples. We mapped the raw reads to the human reference transcriptome. The unmapped reads of each sample were used to analyze the expression levels of these six viruses. We looked at the viral expression throughout all the samples and find out that these six viruses are showing variations (Fig. S2). Further to identify the experiments with very high viral count, we sorted the expression value and used the upper quartile to filter experiments with very high viral level (Fig. S3). We observed that around 100 (10%) samples had very high viral expression count as compared to other samples suggesting the infections of one or more of these viruses (Fig. 2c and 2d) while it does not exclude the possibility that some experiments were infected intentionally for research.

### Identify DE genes involved in viral infection in different cell lines

It is challenging to accurately detect viral counts in RNA-seq data obtained using polyA enrichment chemistry. Therefore, we sought to identify biomarkers that would reveal the impact of the virus on host biology. We analyzed data from two studies (GSE198398 and GSE187420) featuring two different cell lines (Vero 6 and Calu-3) infected with SARS-Cov-2. Using differential expression analysis, we identified 953 and 4101 differentially expressed genes in both studies, respectively. These genes were further analyzed using KEGG analysis to identify pathways related to viral infection, including Influenza A, Herpes simplex virus 1, Hepatitis B, Hepatitis C, and Epstein-Barr virus. We identified 300 differentially expressed genes active in infected samples compared to controls and used these genes in a Random Forest model for feature selection and prediction.

### Biomarker discovery for viral infection in cell lines

After identifying the infected samples in MCF7 cell lines, we collected all (1021 samples) samples normalized expression matrix from ARCH4 (https://maayanlab.cloud/archs4/data.html) and labeled them as infected (110 samples) and control samples (911 samples). To create a gene expression-based biomarker for viral infection, we used a random forest machine learning approach for feature selection and prediction. We used the DE genes from two independent datasets, which were involved in viral infection pathways and active in infection samples as compared to control samples. The training dataset was used for feature selection based on their importance value. We selected the top 12 features (JAK2, ZNF614, ZNF613, PPP2R2A, ZNF595, GADD45B, ZNF433, ITGA5, ZNF627, NFKBIA, PIK3R3, and ZNF333) with an accuracy >0.92 and kappa value >0.44 (Fig. 4a and Fig. 4b). To get more biological insights about these features, we performed the protein-protein interaction, gene ontology (GO) enrichment and KEGG pathways analysis. We observed that only four genes (PIK3R3, NFKBIA, JAK2, and GADD45B) were interacting with each other (Fig. 3c), and all the genes were involved in viral infection, cancer and signaling pathways (Fig. 3d). These features were further used for prediction. Using the random forest to train the features we were able to classify (AUC=0.91) the infected and control samples with an accuracy of 0.93, sensitivity of 0.99, and specificity of 0.60 (Fig. 3e). This suggests that these features could be used to classify infected and non-infected experiments in any cell lines with very high sensitivity. The low specificity means that the biology of few cells infected by viruses was not impacted by the infection.

**Fig. 3.**
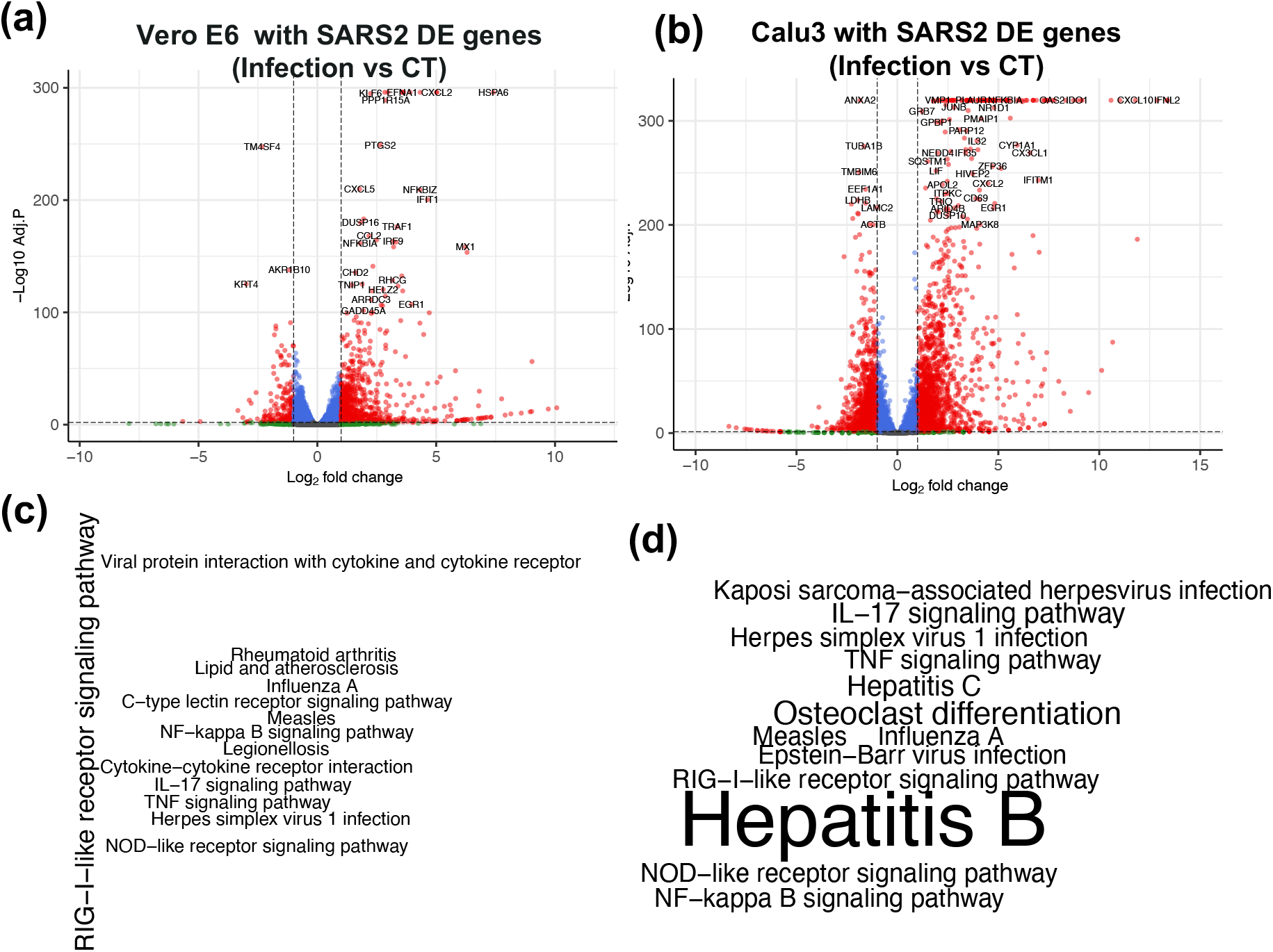
Differentially expressed genes and their KEGG pathways in two different cell lines. Volcano plot showing the differentially expressed (DE) genes in Infection vs Control (CT) experiments with (a) Vero 6 cells and (b) Calu3 cells. (c-d) These genes were used for KEGG pathways analysis to identify the genes involved in viral infection related pathways.

**Fig. 4.**
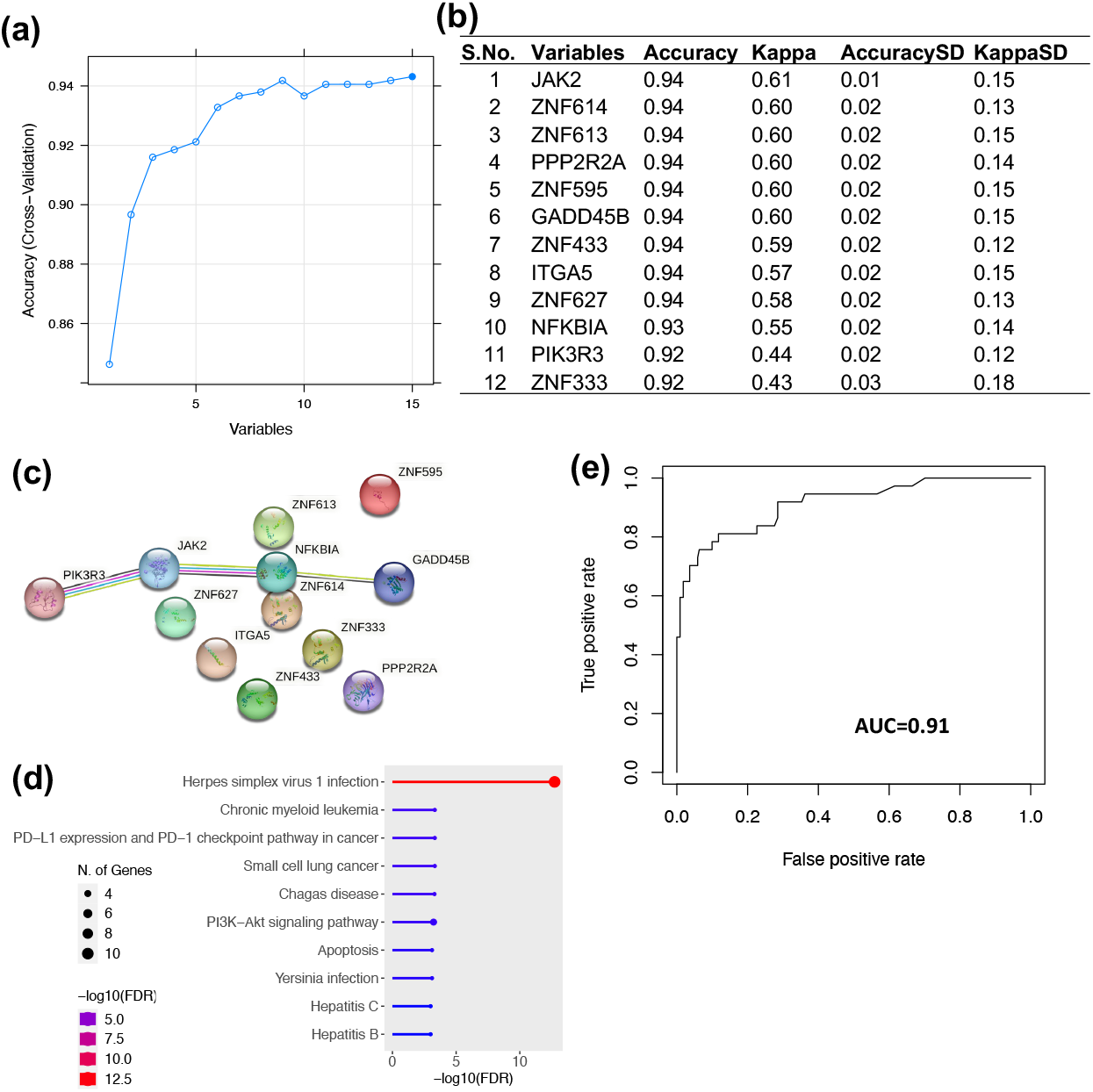
Feature selection and prediction using Random Forest. (a) Line plot showing the top 15 features and their prediction accuracy in the Random Forest. (b) Table listing the top 12 features used for prediction. We selected features with accuracy of more than 0.9, resulting in the removal of three features from the list. (c) Protein-protein interactions of all 12 features were obtained using stringDB. (d) Gene ontology (GO) enrichment of the features identified by Random Forest. (e) Prediction on the test dataset of the MCF7 cell line. An AUC of 0.91, accuracy of 0.93, sensitivity of 0.99, and specificity of 0.60 were achieved.

## DISCUSSION

Cell lines are an important tool in clinical research, allowing scientists to study underlying mechanisms of diseases and develop new treatments. However, there are many challenges when using cell lines for experiments, including the notorious cell contamination. Bacterial and mycoplasma contamination can be mitigated by following best lab practices and often can be easily detected. However, viral contamination is difficult to detect as they cannot be visualized by microscopy. To address this issue, we developed a pipeline called ViralCellDetector to detect viral infections in any sample given its RNA-seq data. ViralCellDetector takes raw sequence reads as input and provides a list of putative viruses. Based on ViralCellDetector, we labeled experiments performed in MCF7 cells as infected and non-infected experiments. Further, we identified the DE genes from two independent datasets belongs to two cell lines and viral infection. The DE genes associated with viral infection were used in machine learning approach to identify the biomarker, which could classify the infected and non-infected cell lines. Using this approach, we identified 12 genes set as biomarkers for viral infection. This biomarker could be used to identify any experiment with viral infection and will help the researchers to take extra measure to avoid any unintentional future contamination in their experiments.

There are many instances of viral infection reported in cell lines (Ali, 2017; Barone *et al*., 2020). Based on their genome type, their mechanism of infection varies. DNA viruses integrate their genome into the host genome and replicate along with cell multiplication, whereas RNA viruses hijack cellular machinery for their own replication. While some viruses, such as HIV, herpes virus, and adenovirus, cause cytopathic effects and can be easily detected, others, such as lactate dehydrogenase virus (LDV), do not cause cytopathic effects and are much harder to detect. In our analysis of MCF7 cell lines, we identified contamination with three DNA viruses and three RNA viruses, including the encephalomyelitis virus, which is an RNA virus and belongs to the herpesviruses family (Griffin, 2011), and the Hepatitis virus which is a single stranded RNA virus and is associated with hepatocellular carcinoma (Liu, 2020), Eptesicus fuscus gammaherpesvirus (EfGHV) is a double stranded DNA virus and mainly infects bat species (Subudhi *et al*., 2018). BeAn 58058 virus is a double stranded DNA virus isolated from the wild rodent Oryzomys species and belongs to the Poxviridae family (Wanzeller *et al*., 2017). Human endogenous retrovirus K113 (HERV-K113) is a type of retrovirus that is found within the human genome, whereas HIV1 is also known to infect humans. Interestingly, all these viruses are used for research in different laboratories, which may be the reason for their infection in cell line experiments.

Further, DE analysis in two independent datasets resulted in a number of DE genes active in infection samples and involved in viral infection. These genes were the great resource for feature selection in machine learning approach. Our machine learning approach resulted in the identification of 12 different genes as biomarkers which were able to classify the infected and non-infected cell lines. Interestingly, all these genes were involved in various viral infection, signaling and cancer related pathways, which showed their importance in viral entry and propagation. Many of these genes are already reported to be supporting the viral infection in the cells (Shen *et al*., 2014; Liu *et al*., 2017; Talledo *et al*., 2012).Their strong association with viral infection and signaling may be the reason for their strong prediction.

Here is some work for future research. Although the biomarker-based approach performs fairly well, we derived the feature set from two datasets. Since previous studies suggested there is a robust host response gene set independent from viruses and their variants (Xing *et al*., 2022; Zheng *et al*., 2021), we will compile more datasets with diverse viruses to refine our biomarkers. Furthermore, our exploratory study suggested the widespread of infection in commonly used cell lines, further work should be performed to uncover the underlying reasons, and further help the community to overcome.

In conclusion, we have developed a computational approach and biomarker to detect viral contamination in cell lines based on readily accessible RNA-seq data. This approach will help researchers understand the negative results of their experiments and take appropriate measures to resolve issues and obtain desirable results. It should be considered as an important aspect of best lab practice guidelines.

## Funding

The research is supported by R01GM134307, and the MSU Global Impact Initiative. The content is solely the responsibility of the authors and does not necessarily represent the official views of sponsors.

## Author Contribution

R.S. conceived and designed the study, performed computational analysis, and wrote the manuscript. S.P. and S.G. help in data analysis. B.C. supervised the study and prepared the manuscript.

## Conflict of Interest

None

## Data availability statement

The source code of ViralCellDetector is freely available at https://github.com/Bin-Chen-Lab/ViralCellDetector/. The data used in this study are public datasets. Those data can be accessible from Gene Omnibus Expression (https://www.ncbi.nlm.nih.gov/geo/browse/?view=series) or Archs4 (https://maayanlab.cloud/archs4/data.html).

---

**Table S1.**
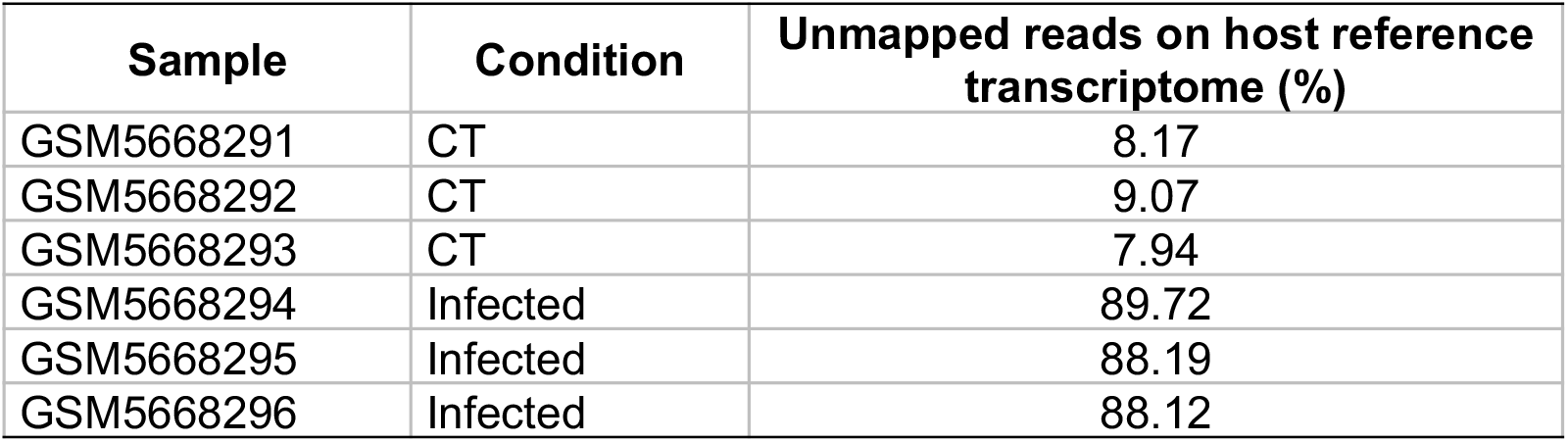
Percentage of raw sequencing reads that were not mapped to the human reference transcriptome.

**Fig. S1.**
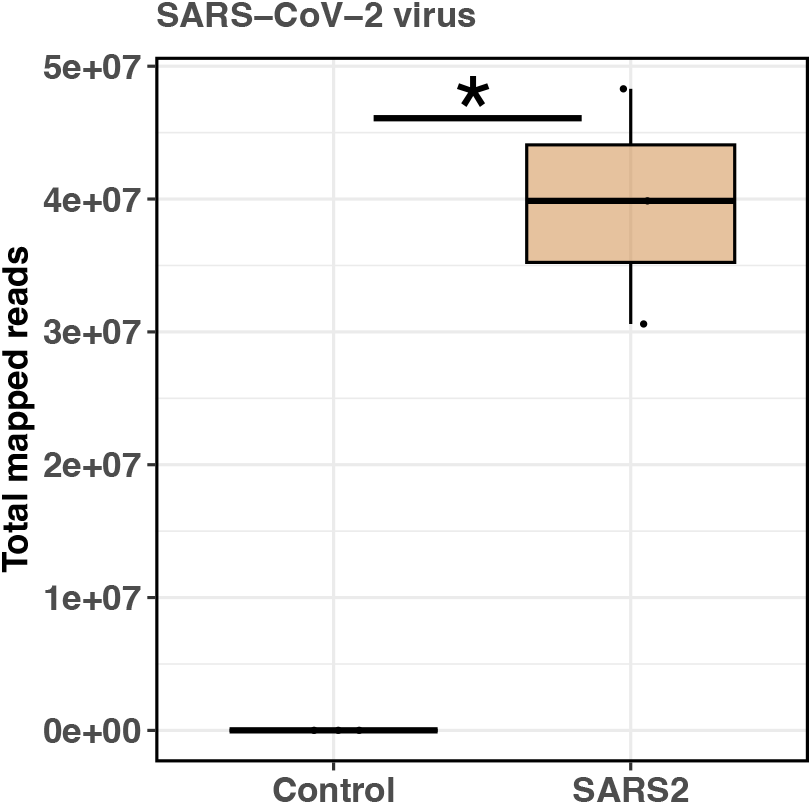
Total mapped reads on the SARS-CoV-2 genome using the VirusCelldetector package. The infected samples showed millions of read mapped on the SARS-CoV-2 genome, whereas control sample has less than 2000 reads belongs to SARS-CoV-2 genome.

**Fig. S2.**
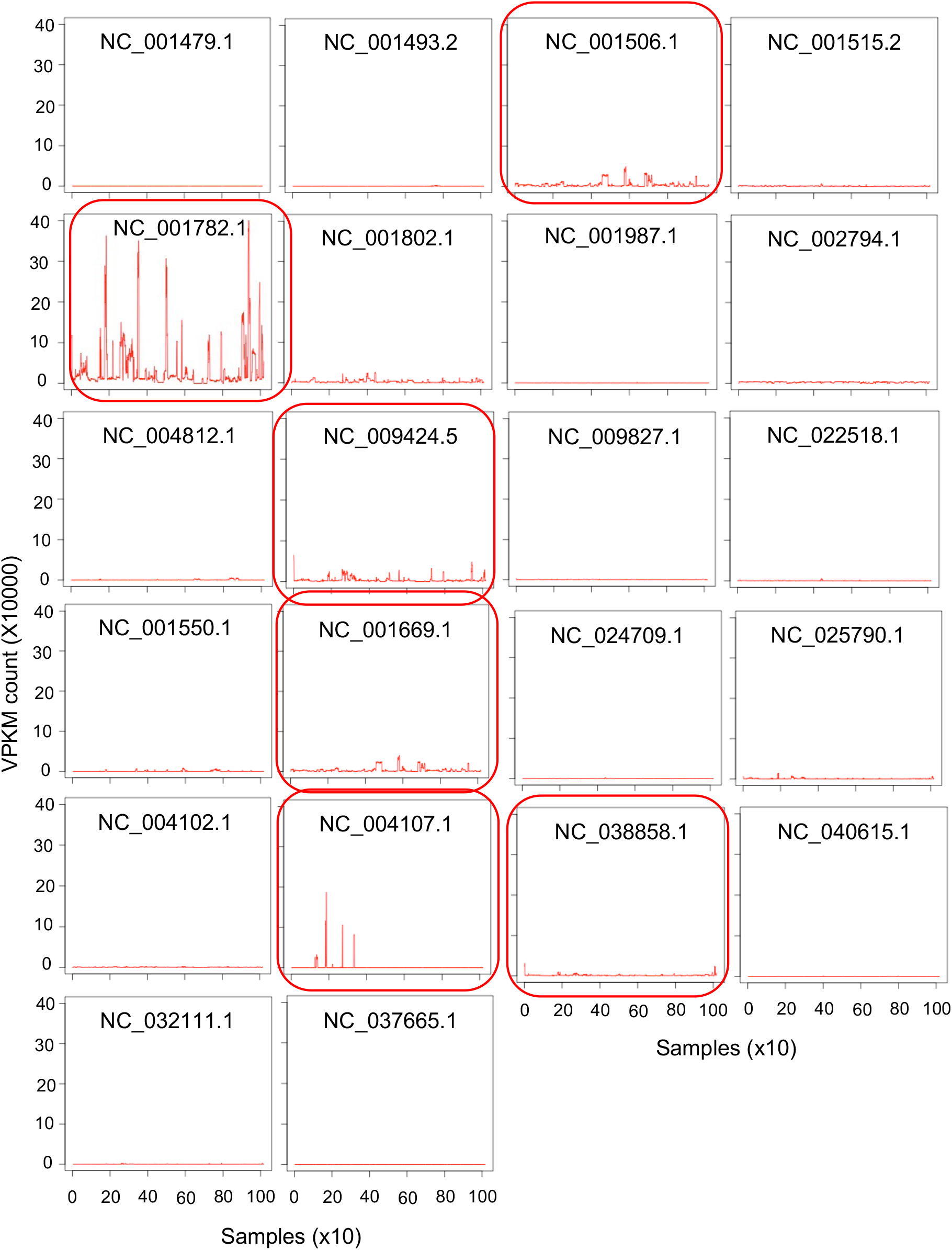
Virus per kb per million reads (VPKM) count of each virus in different experiments in MCF7 cell lines. Only six viruses showed the difference in their VPKM count in different experiments in MCF7 cells.

**Fig. S3.**
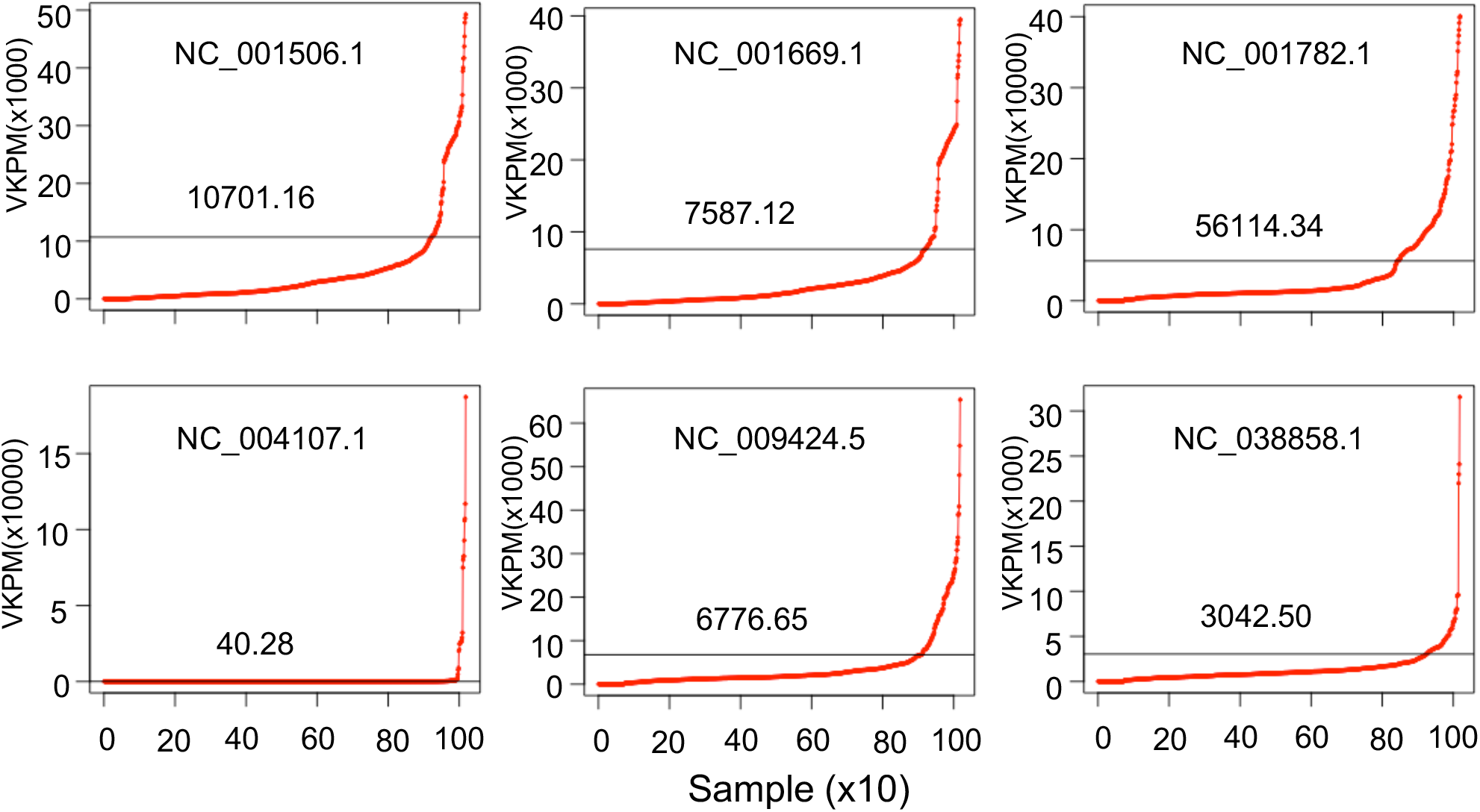
Identification of samples with very high viral expression. We used the upper quartile to filter the samples with very high viral count.

**Table S2.**
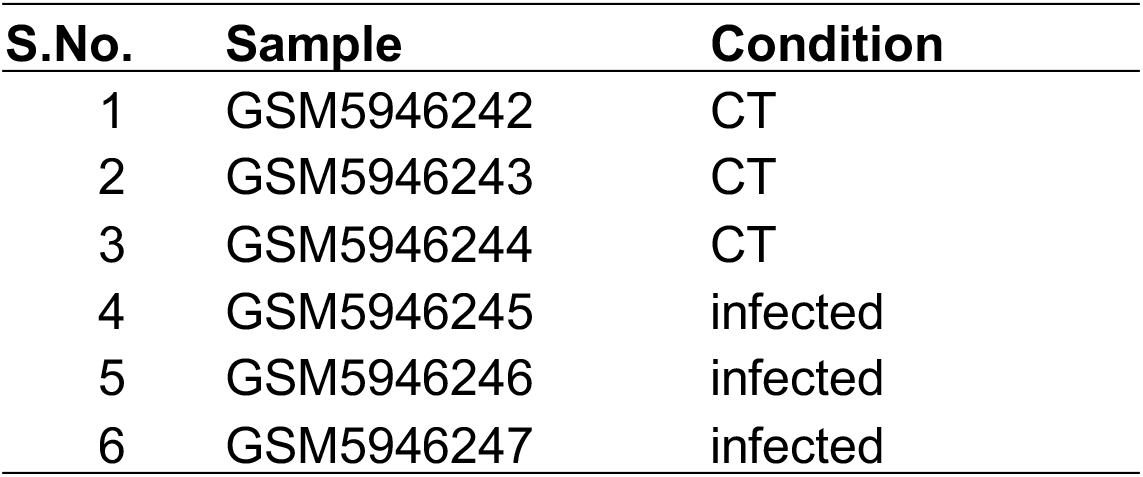
list of samples used for differentially expressed gene analysis in GSE198398.

**Table S3.**
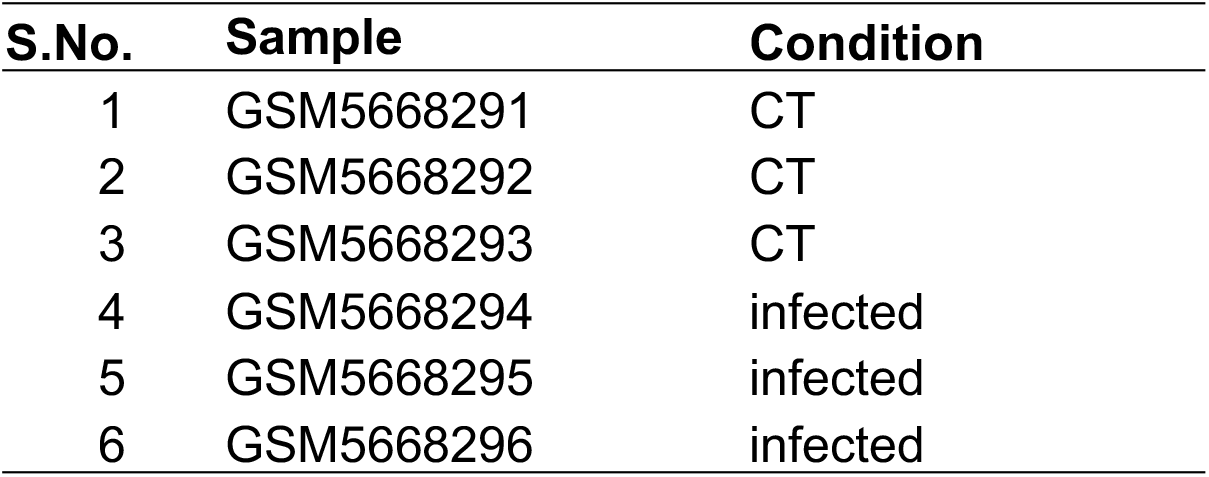
list of samples used for differentially expressed gene analysis in GSE187420.

## Notes

### Competing Interest Statement

The authors have declared no competing interest.

