## Supplement information for "Detection of viral infection in cell lines using ViralCellDetector"

**Table S1.** Percentage of raw sequencing reads that were not mapped to the human reference transcriptome.

| **Sample** | **Condition** | **Unmapped reads on host reference transcriptome (%)** |
|---|---|---|
| GSM5668291 | CT | 8.17 |
| GSM5668292 | CT | 9.07 |
| GSM5668293 | CT | 7.94 |
| GSM5668294 | Infected | 89.72 |
| GSM5668295 | Infected | 88.19 |
| GSM5668296 | Infected | 88.12 |

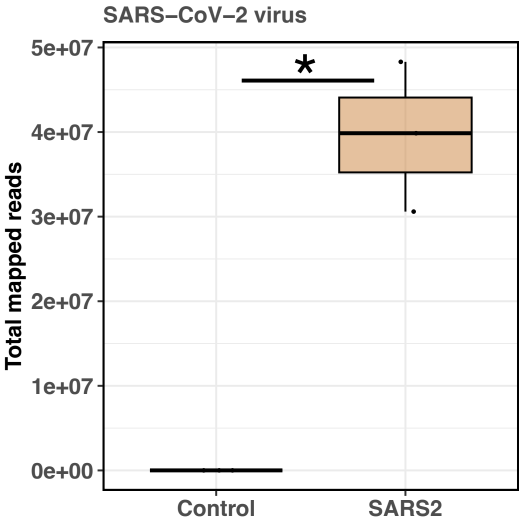


**Fig. S1. Total mapped reads on the SARS-CoV-2 genome using the VirusCelldetector package.** The infected samples showed millions of read mapped on the SARS-CoV-2 genome, whereas control sample has less than 2000 reads belongs to SARS-CoV-2 genome.

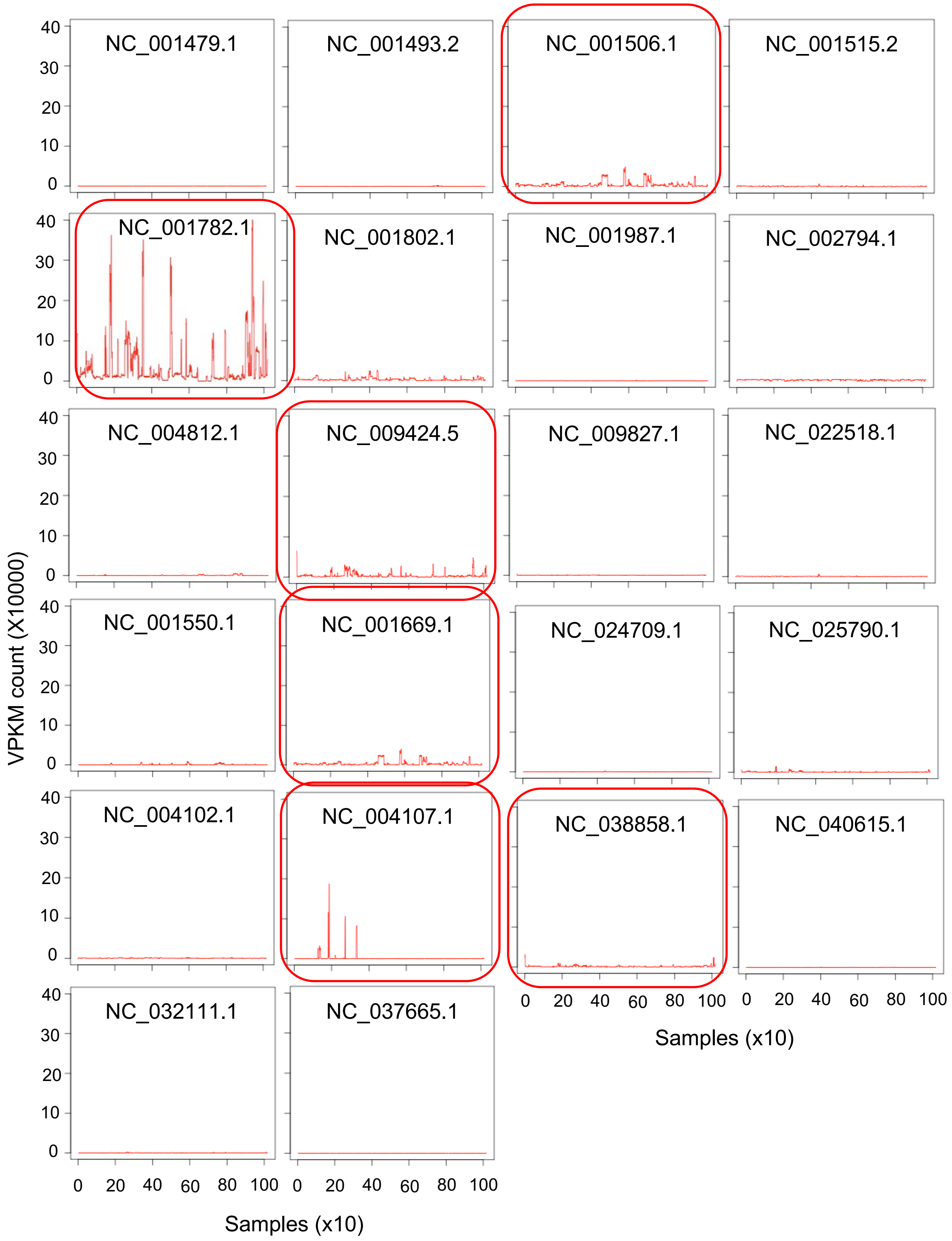


**Fig. S2. Virus per kb per million reads (VPKM) count of each virus in different experiments in MCF7 cell lines.** Only six viruses showed the difference in their VPKM count in different experiments in MCF7 cells.

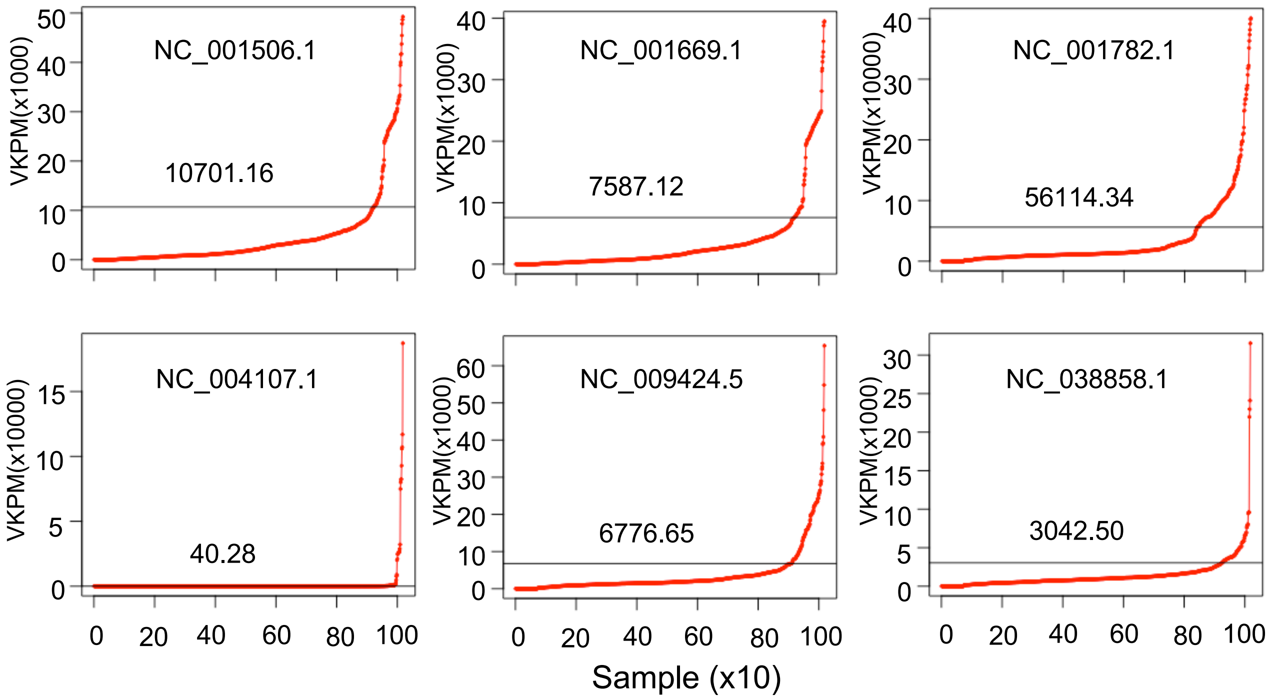


**Fig. S3. Identification of samples with very high viral expression**. We used the upper quartile to filter the samples with very high viral count.

**Table S2** list of samples used for differentially expressed gene analysis in GSE198398.

| S.No. | Sample | Condition |
|---|---|---|
| 1 | GSM5946242 | CT |
| 2 | GSM5946243 | CT |
| 3 | GSM5946244 | CT |
| 4 | GSM5946245 | infected |
| 5 | GSM5946246 | infected |
| 6 | GSM5946247 | infected |

**Table S3** list of samples used for differentially expressed gene analysis in GSE187420.

| S.No. | Sample | Condition |
|---|---|---|
| 1 | GSM5668291 | CT |
| 2 | GSM5668292 | CT |
| 3 | GSM5668293 | CT |
| 4 | GSM5668294 | infected |
| 5 | GSM5668295 | infected |
| 6 | GSM5668296 | infected |
